# Learned response dynamics reflect stimulus timing and encode temporal expectation violations in superficial layers of mouse V1

**DOI:** 10.1101/2024.01.20.576433

**Authors:** Scott G. Knudstrup, Catalina Martinez, Jeffrey P. Gavornik

## Abstract

The ability to recognize ordered event sequences is a fundamental component of sensory cognition and underlies the capacity to generate temporally specific expectations of future events based on previous experience. Various lines of evidence suggest that the primary visual cortex participates in some form of predictive processing, though many details remain ambiguous. Here we use two-photon calcium imaging in layer 2/3 (L2/3) of the mouse primary visual cortex (V1) to study changes in neural activity under a multi-day sequence learning paradigm with respect to prediction error responses, stimulus encoding, and time. We find increased neural activity at the time an expected, but omitted, stimulus would have occurred but no significant prediction error responses following an unexpected stimulus substitution. Sequence representations became sparser and less correlated with training, although these changes had no effect on decoding accuracy of stimulus identity or timing. Additionally, we find that experience modifies the temporal structure of stimulus responses to produce a bias towards predictive stimulus-locked activity. Finally, we observe significant temporal structure during intersequence rest periods that was largely unchanged by training.

## Introduction

Repeated exposure to visual sequences shapes responses in mouse primary visual cortex (V1) in a sequence- and timing-specific manner (Gavornik & Bear, 2014; Price et al., 2023; Sidorov et al., 2020; Tang et al., 2023). While the functional implications of these modifications are unclear, proponents of predictive processing theories have posited that the changes in visually evoked responses over days of spatiotemporal sequence exposure represent a physiological consequence of plasticity through which cortical circuits learn to predict statistical regularities in the environment. According to the predictive coding model (recently reviewed in Keller & Mrsic-Flogel, 2018), heightened responses when visual inputs do not match internally generated predictions constitute a prediction-error signal that carries information about the identity of the unexpected stimulus and the degree of its unlikelihood. While a variety of evidence from primary sensory regions supports this basic framework (Audette & Schneider, 2023; Eliades & Wang, 2008; Fiser et al., 2016; Keller et al., 2012; Stanley & Miall, 2007; Zmarz & Keller, 2016), the degree to which changes of visually evoked responses support predictive coding theories remains a topic of debate. Our lab recently used extracellularly recorded multi-unit activity to investigate how passive exposure to spatiotemporal patterns modifies cells at the layer 4/5 boundary (Price et al., 2023) and found evidence for temporally specific activity consistent with prediction errors in these cells though not to the extent originally observed with LFP-based recordings (Gavornik & Bear, 2014). The convergence of bottom-up and top-down signals in L2/3 suggests that it hosts comparison circuits required in the predictive coding model, an idea supported by recent experiments in the context of visuomotor feedback (Jordan & Keller, 2020).

One notable aspect of our previous work is that responses to a predicted sequence are modified when the constituent elements are presented with the expected order but novel timing. Accurately predicting when events will occur requires forming memories that explicitly encode temporal durations and some sort of internal clock to track elapsed time relative to stimulus events. While temporal processing has been studied in many brain areas, including the hippocampus (Kraus et al., 2013; MacDonald et al., 2011; Pastalkova et al., 2008), entorhinal cortex (Heys & Dombeck, 2018; Kraus et al., 2015; Tsao et al., 2018), sensory thalamus (Komura et al., 2001), and motor cortex (Balasubramaniam et al., 2021) and the visual cortex (Benucci et al., 2009; Price et al., 2023), there are still gaps in understanding how experience encodes temporal expectations into neural circuits.

To investigate the role superficial cortical layers play in coding time and temporal predictions, we used an implicit sequence learning paradigm and two-photon calcium imaging to study how exposure to visual sequences shapes L2/3 neuron responses and how responses are modified when ordinal or temporal expectations are violated. After baseline imaging sessions, mice were exposed to a single training sequence for four consecutive days. On the fifth day, we exposed the mice to the training sequence and two test sequences designed to elicit temporal or ordinal prediction errors. We found little evidence of prediction errors when elements were presented with an unexpected order but did find elevated activity at the time an omitted element should have been presented which we interpret as a form of temporal prediction error. Though evoked response dynamics are modified with training, decorrelating and shifting towards stimulus-locked responses, this has no obvious advantage for decoding elapsed time or stimulus identity at the population level. Finally, we find temporally specific activity in the gray inter-sequence period.

## Results

### Experimental design

We used a multiday experimental protocol in which mice were exposed to a standard sequence (ABCD) and two variants (ABBD and ACBD) designed to elicit positive (unexpected element C in the B position) and negative (element C omitted by keeping B on the screen for twice the expected duration) prediction errors (figure 1A), where the terms “positive” and “negative” are used to distinguish between cases with unexpected stimuli vs the lack of an expected stimuli. To establish a baseline for comparison, mice viewed all three sequences on day 0 (pre-training).

**Figure 1:**
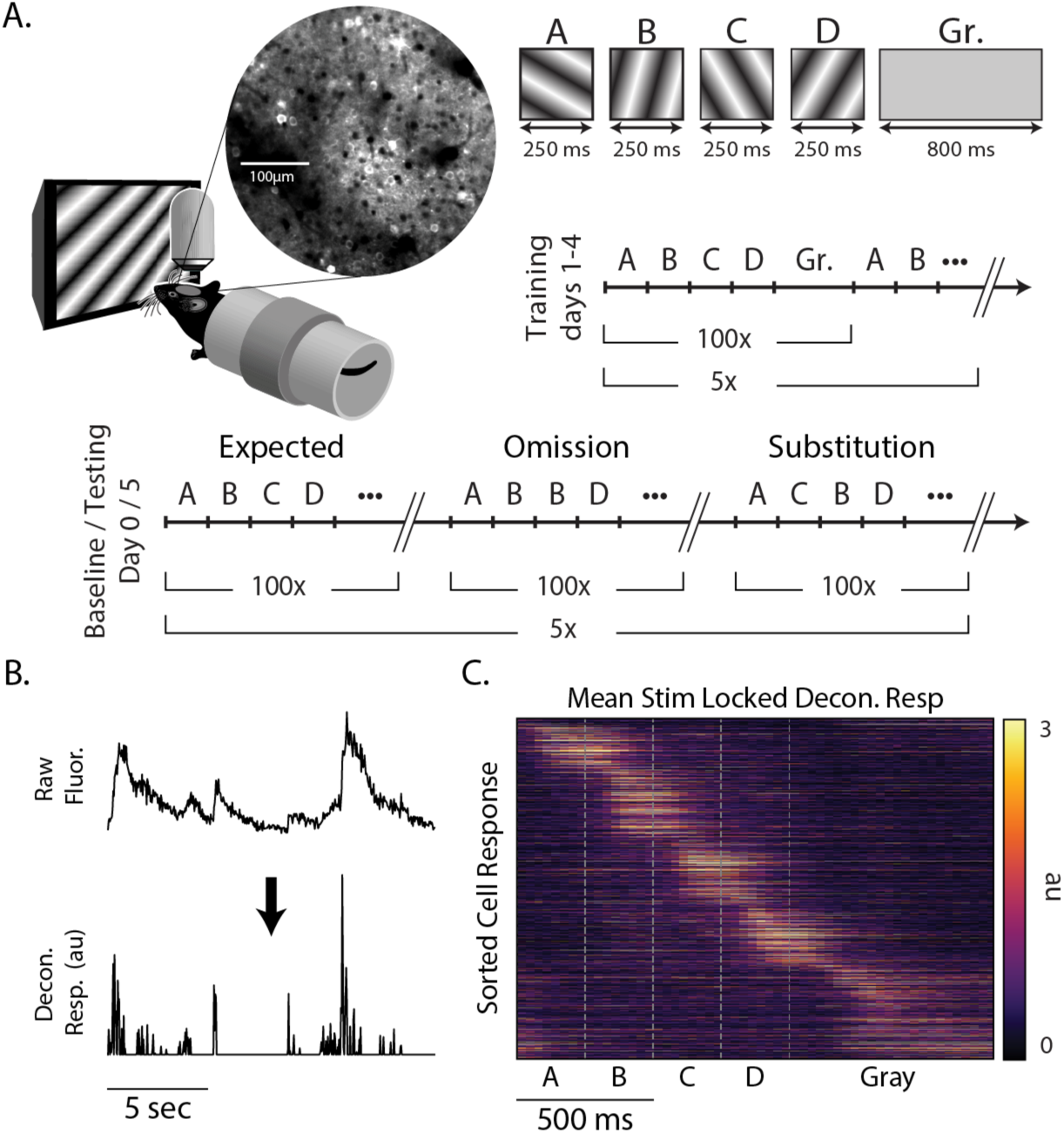
**Experimental design** (A) Awake head-fixed mice viewed sequences of oriented gratings while undergoing two-photon calcium imaging in superficial V1, the location of which was determined using widefield retinotopic mapping. To establish a functional baseline, mice were shown the sequences ABCD, ABBD, and ACBD on day 0 (pre-training). After seeing only ABCD during training (days 1-4), mice were again shown all three sequences on the test day (day 5). Sequence elements were held on the screen for 250 ms each and presented without gaps between elements. Sequence presentations were separated by 800 ms of gray screen. (B) To increase the temporal specificity of calcium signals, fluorescence extracted from ROIs marking individual neurons was deconvolved prior to analysis. (C) Heat-map showing the averaged responses of 1368 cells on day 0 to sequence ABCD. To remove artificial temporal structure, this map represents that average activity of odd trials sorted based on the time of peak activity in even trials.

After two days with no visual stimulation, mice were then shown ABCD exclusively during days 1-4 (training). On day 5, mice saw all three sequences again (testing). The training period serves to build an expectation for the sequence ABCD, while the day 0 baseline allowed us to compare responses to all sequences before and after ABCD had been established as the expected sequence.

Mice were awake and head fixed in all sessions. Sequences were composed of four isoluminant oriented gratings, each of which was presented for 250 ms for a total sequence length of 1 second, and sequence presentations were separated by 800 ms of gray screen. Sequences were presented in 5 blocks of 100 with a 10 second gray period between blocks. Since gratings transitioned directly into one another with no gap, element B in ABBD was functionally a single element lasting 500 ms with no visual indicator of when the first B ended and second began. We notate individual sequence elements using bold lettering (e.g., A**B**CD refers to B in ABCD). As detailed in the Methods section, the average pixel fluorescence value within ROIs designating visually responsive somas was calculated for each frame. This signal was deconvolved in time (Pachitariu et al., 2016) to produce an activity metric with a relatively high degree of temporal precision compared to the slow underlying calcium signal (figure 1B). This approach produced a population of neural responses that clearly shows unique responses to each element of the sequence with activity spanning the entire period of stimulation (figure 1C).

Binocular visual cortex was identified via retinotopic stimulation, and landmarks from reference images taken on day 0 were used to target the same approximate population of neurons on day 5 (supplementary figure 1). We imaged a total of 1368 and 1500 cells on days 0 and 5, respectively, from 8 mice. Cells were categorized by stimulus selectivity and visual responsiveness within the sequence (see Table 1). We did not track the response properties of individual neurons across training days, but the percent of neurons representing each element decreased by an average of 2.8 % between days 0 and 5, while the number of cells classified as gray responsive stayed approximately the same.

**Table 1:** Stimulus selectivity Stimulus selectivity for neurons recorded on day 0 (n=1368) and day 5 (n=1500). A cell was considered stimulus-selective for an element if the average activity evoked by that stimulus was more than two standard deviations higher than any other stimulus.

| Stimulus | Day 0 | Day 5 | % change |
| --- | --- | --- | --- |
| A | 84 (6.1 %) | 41 (2.7%) | -3.4% |
| B | 137 (10.0%) | 107 (7.1%) | -2.9% |
| C | 84 (6.1 %) | 39 (2.7%) | -3.4% |
| D | 126 (9.2%) | 65 (4.3%) | -4.9% |
| Gray | 129 (9.4%) | 154 (10.3%) | 0.9% |

### Omissions, but not substitutions, drive prediction errors

The first goal of our experiments was to test whether layer 2/3 neurons display prediction errors in response to stimulus omissions or substitutions. The sequence ABBD contains an omission violation in position 3 when B is held on screen. The sequence ACBD contains a substitution violation at the second element since C appears where B is expected. The predictive coding model holds that both forms of expectation violation should result in elevated responses on day 5 relative to the day 0 baseline as diagramed in figure 2A.

**Figure 2:**
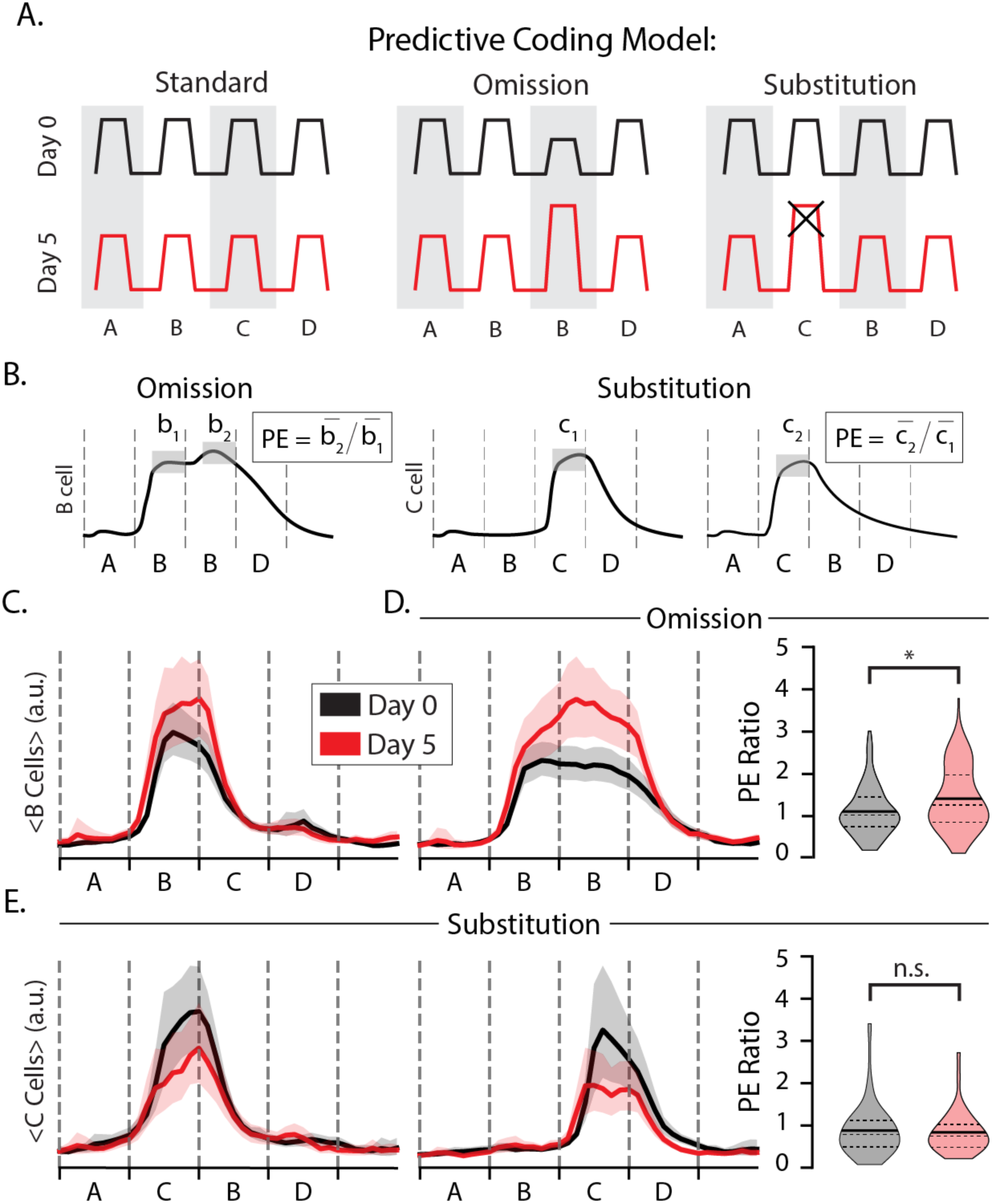
**Stimulus omissions drive prediction errors** (A) Diagram of putative prediction error (PE) responses to omissions (middle) and substitutions (right) where the omitted/substituted element is expected to drive elevated responses on day 5 compared with day 0 (the x indicates that this elevated response is not present in our data). (B) PE ratios were computed by dividing trial- and time-averaged activity during the deviant image by activity during a corresponding standard image (traces in panel B were drawn manually for illustration purposes). (C) Average trace of B-responsive cells to ABCD with bootstrapped 95% confidence intervals on days 0 (gray) and 5 (red). (D) Average trace of B-responsive cells to ABBD (left) and distributions of omission-type PE ratios. Note that there is no change in visual stimulus at the B_1_B_2_ transition. On day 0, mean PE=1.10 (n=137). On day 5, mean PE=1.44 (n=107). Distributions were significantly different (p << 0.05; n=244; KS-test). (E) Average trace of C-responsive cells to ACBD (left) and ABCD (middle). (right) Distributions of substitution-type PE ratios for C-responsive cells. On day 0, mean PE=0.88 (n=84). On day 5, mean PE=0.84 (n=39). Day 0 and day 5 distributions were not significantly different (p > 0.05; n=123; KS-test).

We first classified cells based on their stimulus selectivity with and without regard for sequence context, labeling cells as selective for a specific element when the average activity evoked by that element was two standard deviations higher than activity evoked by other stimuli. This approach was designed to identify cells that, for example, fire preferentially to element C in at either ABCD or ACBD. In this example, a cell that fires more during A**C**BD than AB**C**D would exhibit a pattern of excessive activity associated with prediction error signals. We quantified this effect using a prediction error metric (PE) where the trial- and time-averaged activity during an unexpected event is divided by the activity during a similar but expected event (figure 2B). If training drives changes to prediction error responses, then the distribution of PE ratios should be significantly different after training on day 5 than in baseline day 0 recordings.

To look for omission errors, we identified B-responsive cells and computed their PE ratios using AB**B**D and A**B**BD as the unexpected and expected stimuli, respectively (figure 2B, left). The average response to element B in position 2 increased slightly with training (figure 2C) and was approximately the same during the second position regardless of which stimulus followed on days 0 and 5 (figure 2 C, D). Training produced significant differences in the temporal characteristics of B-responsive cells. On day 0, activity in B-responsive neurons decreased consistently as the B element was held into position 3. After training, however, activity was elevated during this period with a slight increase following the point in time at which element C would normally be seen (figure 2D). The PE ratio increased significantly with training (p = 0.005, n=244, KS-test). We interpret this result, wherein evoked activity increases at a point in time where visual inputs are unchanging, as representing a temporally specific prediction error following the omission of an expected visual transition. This finding is broadly consistent with our previous findings in deep layer 4 (Price et al., 2023) and suggests that temporal prediction errors can be found in the population of excitatory neurons selective for an expected stimulus. We repeated this analysis of PE ratios using A**B**CD as the expected/standard element and found the results to be generally insensitive to the choice of A**B**CD vs A**B**BD (supplementary figure 2). We did not clearly identify a separate population of otherwise sequence non-responsive “prediction error cells” uniquely activated during stimulus omissions as posited by some predictive coding models (supplementary figure 3). Out of the 107 B-responsive cells on day 5, only one appeared to fire exclusively during the omitted stimulus.

To look for substitution errors, we identified C-responsive cells and computed their PE ratios using the C response in both the unexpected (A**C**BD) and expected (AB**C**D) positions (figure 2B, right). If substitution drives a prediction error following an unexpected substitution, the PE ratios for these cells would be approximately equal to 1 at baseline and higher than 1 after training. The average response to element C was higher at baseline than on day 5 (figure 2E). We found that mean PE ratios were approximately equal to 1 on both days, with a slight but statistically insignificant decrease following training (p = 0.64, n = 123; KS-test). Contrary to our expectations coming into this experiment, training did not facilitate a significant change in responses to the expected vs unexpected element during element substitutions. As with the omission case, we did not identify a unique population of otherwise visually non-responsive “prediction error cells” following substitution (supplementary figure 4). Overall, and in contrast to our previous results from deeper layers, we find no evidence that unexpected ordinal substitutions drive elevated activity consistent with predictive coding models in excitatory layer 2/3 cells. Given the hierarchical structure of the data, we also performed an analysis on omission- and substitution-type prediction errors using the non-parametric hierarchical bootstrap to estimate mean traces and PE ratios, the results of which are consistent with the analysis presented here (supplementary figure 8).

### Experience modifies activity in principal component space and drives sparsification

Sequence responses on day 5 look qualitatively different at the population level compared to baseline, losing the blocky alignment with element transitions of stimulus-selective populations (figure 3A). To quantify this observation, we used principal component analysis to identify the form of latent variables underlying population dynamics in pre- and post-training datasets. Prior to training, stimuli drive activity along several axes in principal component space in complex combinations, and the principal components do not neatly reflect activity driven by any particular sequence element (figure 3B). In contrast, activity in principal component space is discretized or “untangled” after training with clearly defined peaks reflecting sequence elements. Though the total number of principal components required to account for 90% of total variance did not change appreciably with training (approximately 10 PCs on days 0 and 5), the components accounting for the majority of variance shifted to neatly reflect the specific sequence structure after training.

**Figure 3:**
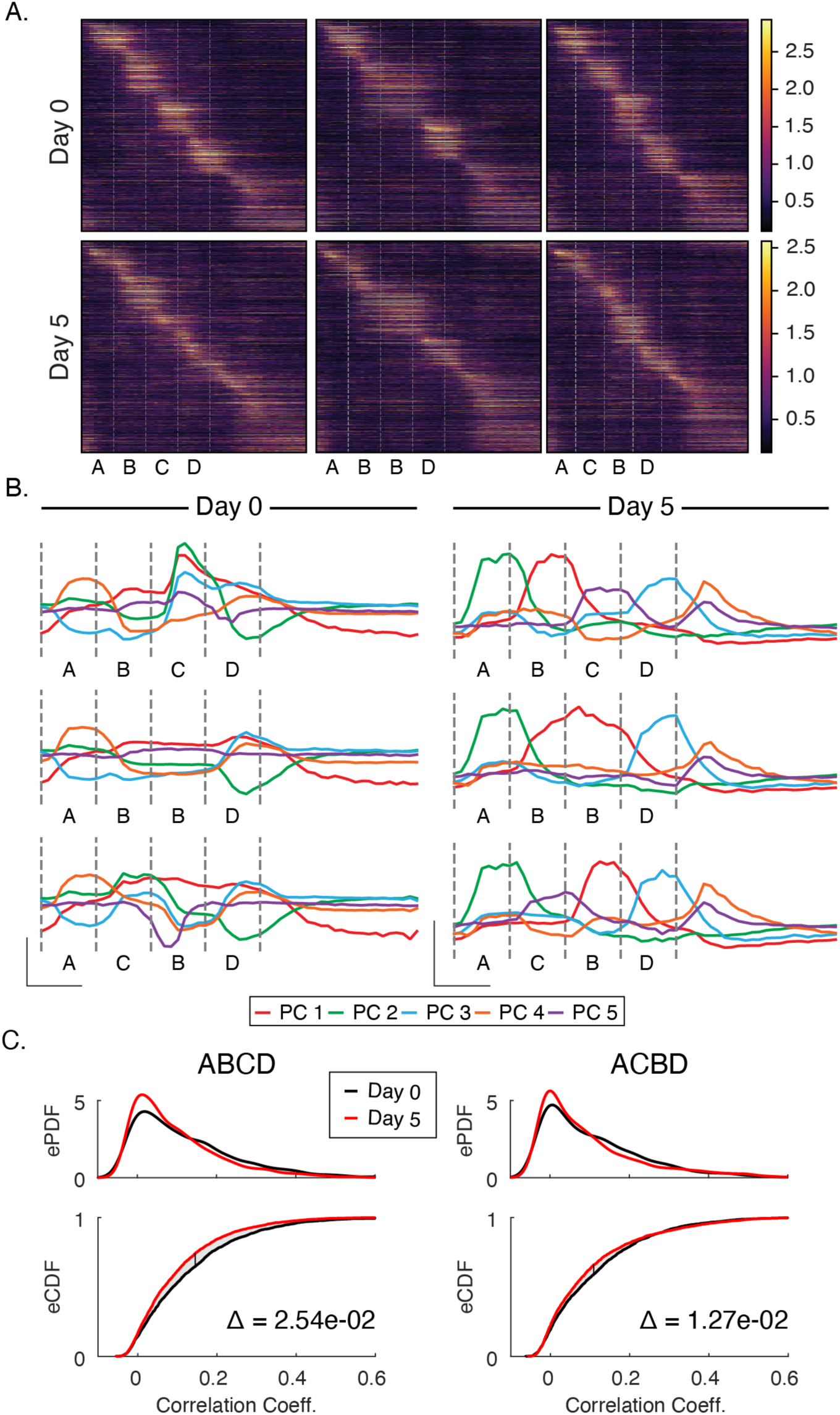
**Principal Component and Sparsity Analysis** (A) Trial average population responses, sorted by time to peak latency in each cell, to each sequence before and after training. (B) Prior to training, activity is driven along principal components jointly in complex combinations. After training, the most significant principal components align neatly with individual stimuli. In both datasets, the first five components explain ∼80% of the variance. (C) To test whether changes in principal component space reflected the decorrelation of responses, we computed Pearson-correlation coefficients between all four images for each sequence presentation individually. Empirical PDFs (top panels) and CDFs (bottom panels) of Pearson-correlation coefficients. After training, activity became significantly less correlated (p < 0.05; n=6000; KS-test) for ABCD and ACBD. Delta (Δ) on bottom panels indicates area between curves on CDFs.

Training also facilitated a reduction in the fraction of neurons that responded preferentially to individual stimuli. Prior to training, 40.8% of neurons were driven more than two standard deviations above their baseline firing rate by one of the stimuli compared with 27.1% after training. We also quantified how many cells were “visually modulated”, a more permissive metric that simply compares activity between gray and non-gray periods. Under this measure, the percentages of visually modulated cells went from 72% on day 0 to 60% after training. As measured by the fraction of cells participating in visually modulated or stimulus-specific activity, responses are sparser after training than at baseline.

Since information theory holds that codes gain efficiency by eliminating redundant information (see Price & Gavornik 2022 for how this relates to predictive coding in the visual system), we reasoned that correlations between stimuli responses would decrease with training. To test this, we examined correlation coefficients between stimuli to determine activity on day 5 was less correlated than on day 0. For each sequence presentation, we calculated the Pearson correlation coefficients between all pairs of stimuli within a sequence (A vs B, B vs C, etc. see Methods), and plotted empirical distributions of these coefficients before and after training (figure 3C). We found that correlations were reduced by 10-20% on day 5 compared with day 0. Across all sequences, the shift in correlation distributions was significantly different between days (ABCD: p < 1e-3; n=6000. ACBD: p < 1e-3, n=6000; KS-test). This, coupled with the overall decrease in number of cells responding to visual stimulation, suggests that experience-dependent spatiotemporal plasticity increases coding efficiency.

### Plasticity does not increase decoding accuracy

Our principal component analysis suggests that experience creates representations that are more easily separable in high-dimensional space. To test whether experiential shaping of cortical circuits increases the ability to differentiate between different visual stimuli, we trained a linear decoder with responses from all stimulus contexts (figure 4A). We found that accuracy was well above chance (1/15 = 6.7% ± 0.4%) on day 0 (78%) and day 5 (76%) and that the decoder was able to discriminate between cases we expected to be indistinguishable. For example, we expected **A**BCD would be more-or-less indistinguishable from **A**BBD and **A**CBD since A occurs first in each sequence and is always preceded by a long (800 ms) gray period. This was not the case. Most often, the decoder correctly identified which sequence stimulus A came from. This pattern was observed with other cases as well, including the intersequence gray periods. This prompted us to consider whether responses undergo long-timescale alterations over the 30 min session that led to separability in high-dimensional space.

**Figure 4:**
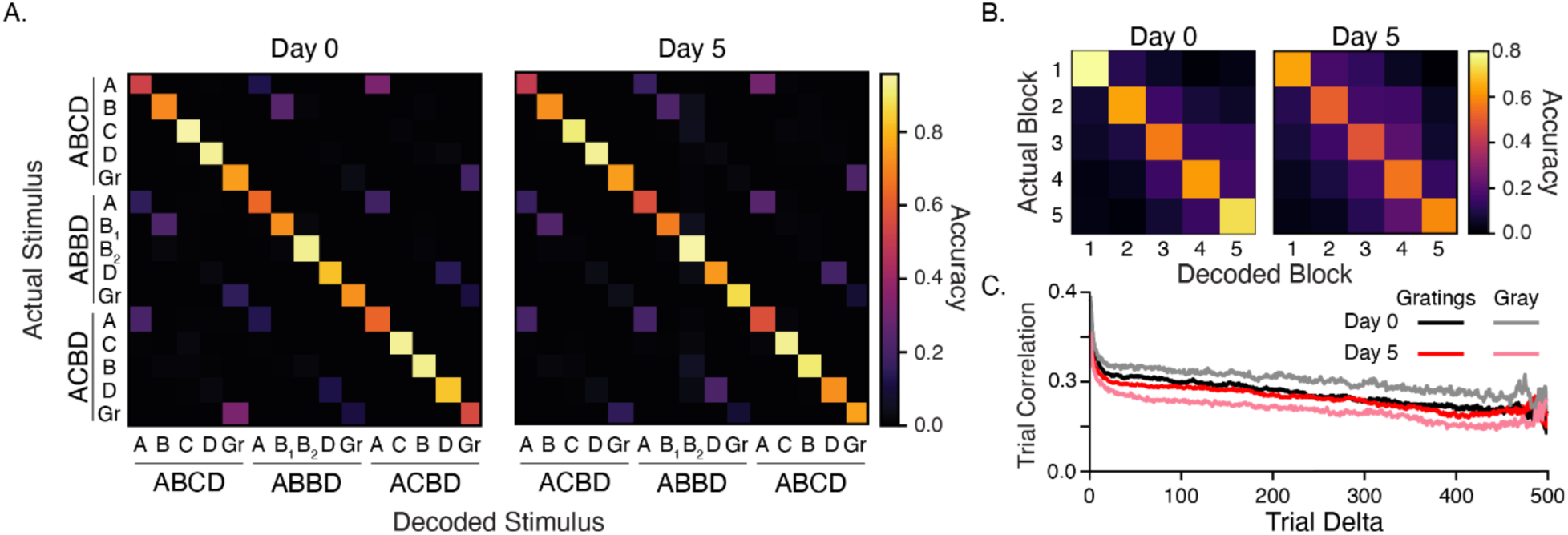
**Stimulus decoding and representational drift** (A) Average confusion matrices (100 iterations) for decoders trained on responses to all images. Average decoder accuracy was 78% and 76% for days 0 and 5, respectively. Both are well above chance (6.7% ± 0.4%). The ability to differentiate correctly between the same image in different contexts, such as **A**BCD vs **A**BBD, prompted us to consider whether responses slowly drifted over time since sequences were presented in blocks of 100. (B) A decoder trained on individual elements (e.g., **A**BCD) accurately classifies which block responses came from, with errors decreasing along with distance between blocks (e.g., block 2 responses are more often classified as belonging to block 3 than blocks 4 or 5). Confusion matrices represent averages over decoders trained on stimuli individually (see Methods). Decoder accuracy was 68% on day 0 and 56% on day 5. (C) We estimated drift by computing Pearson’s correlation coefficients between all pairs of population vectors driven by a particular sequence element and grouped these values based of between pairs were in time/trial. Responses become less correlated as distance between trials increases during both stimulus-evoked and gray periods. The largest change in overall temporal correlation was seen between gray periods on days 0 and 5.

We next trained a decoder with data from the same sequence element (e.g., **A**BCD) and found that the decoder accurately predicted which block the stimuli came from, and that the rate of false positives decreased with the distance between actual and predicted blocks (figure 4B).

This suggests that there is a slow, continuous drift in representations that gradually separates responses in vector space over the course of a single imaging session. In principle, this could be due to photobleaching, repetition suppression, adaptation, or slow changes in arousal that exert a rescaling effect on neural responses. Repeating the same analysis using normalized vectors to train the decoder did not degrade the decoder’s ability to accurately predict which block a response came from (supplementary figure 5).

To further characterize slow changes to representations over time, we performed an analysis of vector similarity as a function of the distance between response pairs to the same stimulus (figure 4C). For each stimulus, we constructed a population vector for each trial by time-averaging data over the presentation period. We then calculated the Pearson’s correlation between all pairs of population vectors and grouped correlation coefficients by the distance (in time/trial) between pairs. The average correlation values for each interval capture the similarity between representations. Calculating mean correlations as a function of the interval between trials showed a gradual decline in vector similarity, indicating drift across the 30-minute recording session. We repeated this analysis for animals individually and found the trend towards decreasing correlations held in individual mice (supplementary figure 6). This finding is consistent with previous work in V1 that used a similar metric to quantify representational drift (Deitch et al., 2021).

### Experience facilitates stimulus-locked responses

To understand how evoked dynamics shift with training, we assigned a time bin to each cell based on its point of maximal activity (figure 5A). At baseline, few cells have peak activity times immediately after stimulus onset, and the percent of cells with late maximal response times increases gradually over each stimulus (figure 5B). After training, the pattern is reversed. More cells are active at or near stimulus onset or immediately before the next element transition, while the number of cells with peak activity at intermediate time points between element transitions decreases. There is no obvious shift in response latencies following stimulus substitution or omission. These shifts comport with the changes discussed in figure 3 and could represent temporal representations being learned at the population level. To test this idea, we trained a linear decoder to report time within a stimulation period. The decoder was able to report time above chance within the stimulation period for all sequence in both naïve and trained networks (figure 5C).

**Figure 5:**
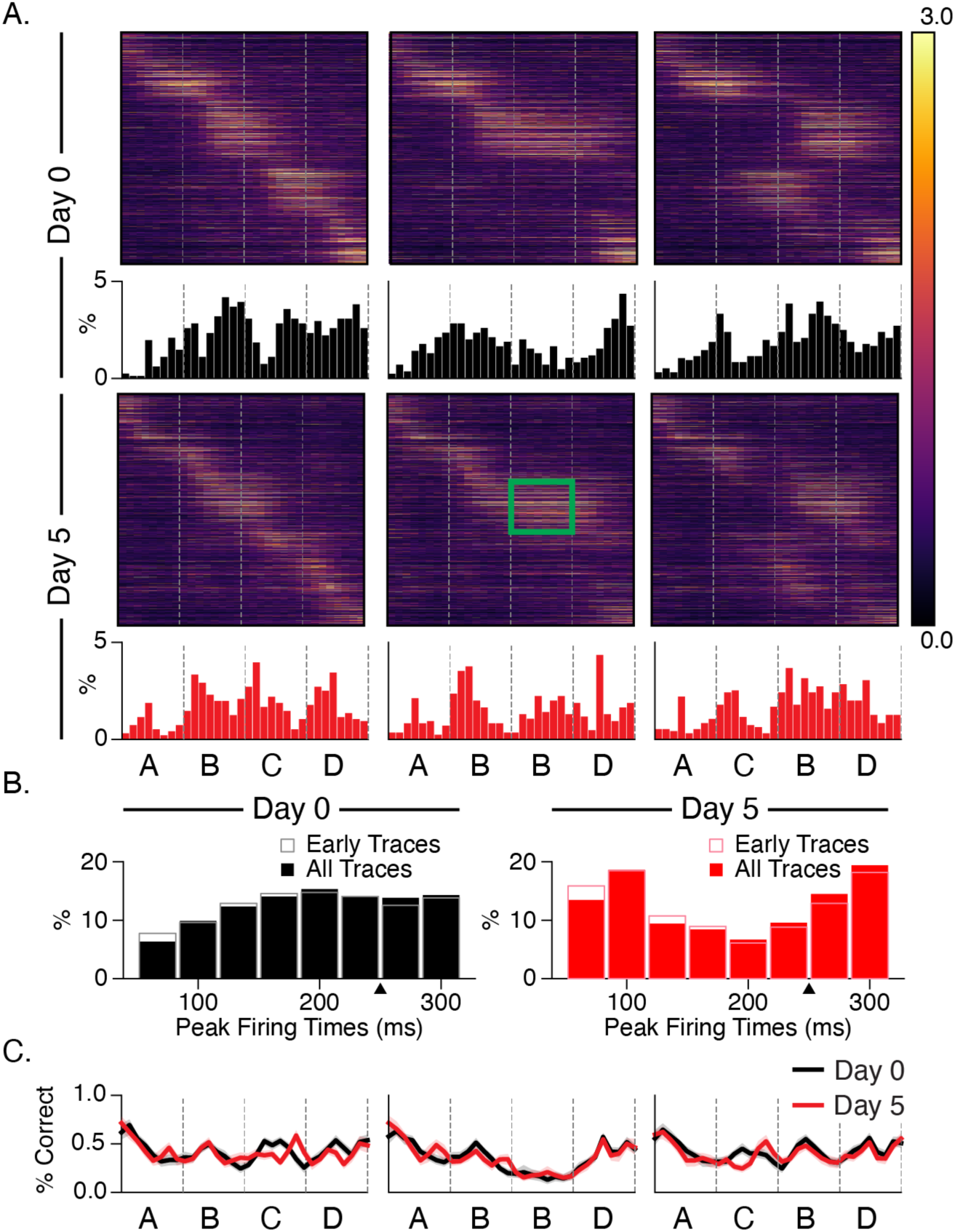
**Stimulus responses shift with training** (A) Heatmaps displaying trial-averaged responses to all sequences sorted by peak response time during ABCD. Histograms showing the binned location of peak activity for each cell shift towards stimulus-locked phasic responses after training. The green rectangle highlights the location of increased activity on day 5 relative to baseline. (B) Combining histograms across all stimulus conditions shows how temporal response latency patterning changes over days. Note that the first two time bins (66 ms) after onsets are omitted in histograms to compensate for the transmission delay from retina to L2/3 cells. Prior to training, responses slowly build up after onset. After training, responses are robust at onset and undergo quick depression prior to ramping up for the next element. Histograms based only on early trials (first 100) show that this pattern does not change significantly over the course of the imaging session. (C) To test whether changes to temporal patterning might reflect a change in the ability to discern durations, we trained decoders on responses from different time bins following sequence onset. Decoder accuracy did not change significantly as a function of training or sequence.

To determine how training changes activity between sequences, we next performed the same analysis on the intersequence gray periods and found that a subpopulation of cells was activated ∼300 ms after gray period onset (figure 6), slightly later than the 250 ms interval suggested by the temporal structure of our visual sequence. A second population of cells exhibited ramping activity leading up to the end of the 800 ms gray period (supplementary figure 7). While this response looks like it anticipates the onset of the next sequence, it is present at baseline, including in early trials, and did not change significantly with training. The only noticeable difference after training is a decrease in the percentage of cells firing early in the gray period on day 5 relative to baseline.

**Figure 6:**
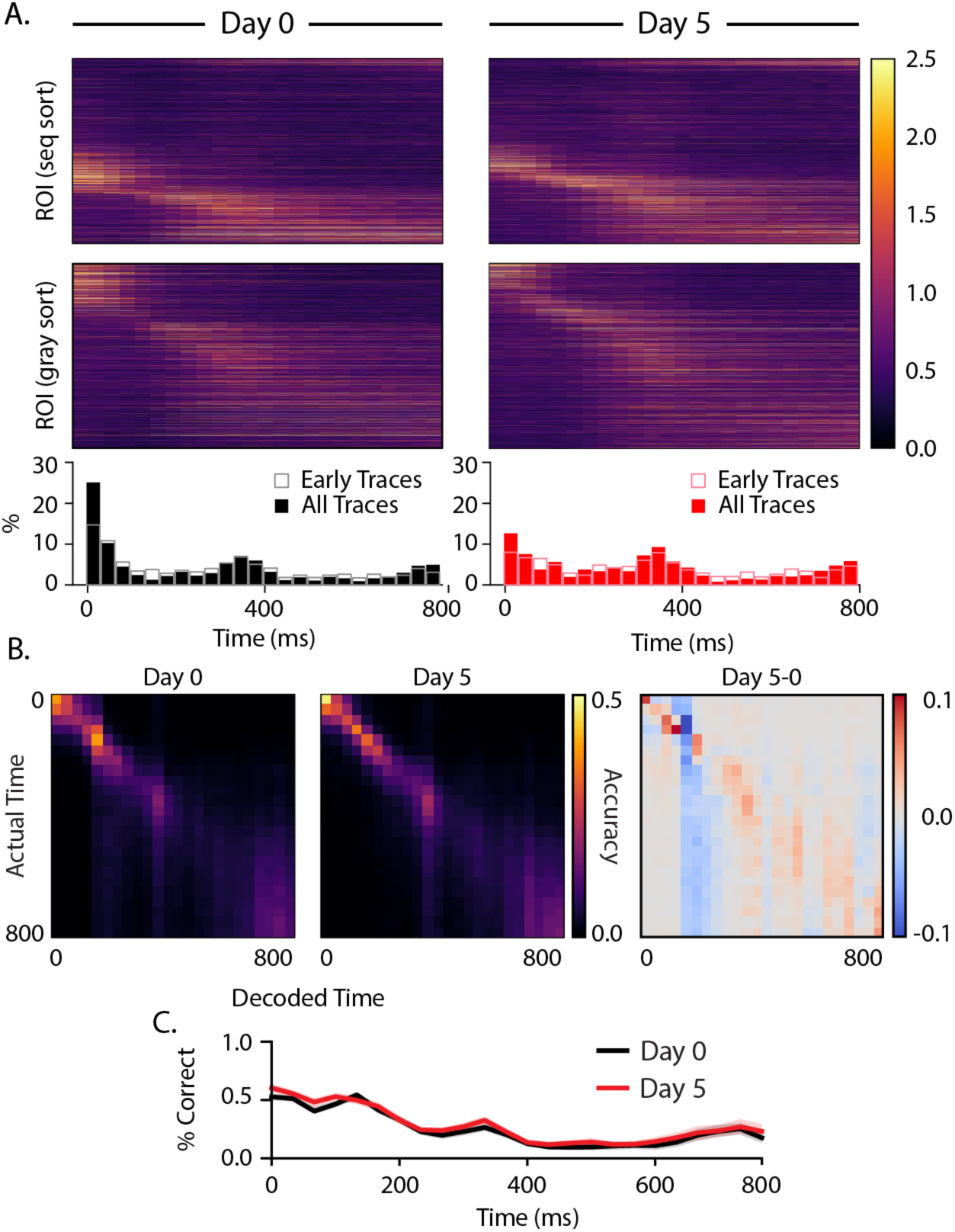
**Dynamics during intersequence gray periods** (A) Heatmaps displaying trial-averaged responses during gray periods following sequence presentations. Activity is shown sorted as before by peak response time during ABCD presentations (top panels) and resorted based on peak response during the gray periods only (middle panels). Histograms (bottom) reflect locations of peak activity for each cell and show that there is an increase in active population size about 300 ms after gray onset and a second uptick towards preceding the next sequence presentation at 800 ms. (B) Average confusion matrices (100 iterations) for decoders trained on responses at different delays from gray onset for day 0 (left), day 5 (middle), and the difference between them (right). (C) Overall decoder accuracy as a function of time since gray onset did not change significantly with training.

As before, we trained a decoder on peak activity time bin population vectors asked the decoder to classify testing data by which time bin it belonged to. While time decoding was slightly more accurate on average along the confusion matrix diagonal (Fig. 6B), there was no significant change in temporal decoding during intersequence gray periods (Fig. 6C). On average, decoding accuracy was 23.4% on day 0 and 23.4% on day 5. In both cases, the decoder performed well above chance (3.1% ± 0.2%). There are moderate increases in decoding accuracy in about 300-400 ms and nearing the end of the 800 ms gray period coinciding with activity peaks at these times. As was the case with responses evoked during sequence presentation, neural dynamics are sufficient for a linear decoder to determine time fairly accurately within the gray period before and after training.

## Discussion

In this study, we used chronic two-photon imaging to measure activity in L2/3 of mouse V1 during passive exposure to standard and deviant visual sequences before and after a four-day training period. This study was designed to test the hypotheses that neurons in L2/3 exhibit prediction error responses elicited by expectation violations by comparing how V1 responded to identical sequences before and after exposure established an expectation of how different sequence elements relate to each other in time. We focused on two types of violations (omissions and substitutions) and found mixed support for hypothesized prediction errors. We also found that experience yielded a sparsification of neural responses characterized by a decrease in the percent visually responsive cells and a decorrelation of stimulus responses. Response vectors in principal component space were altered by experience, yielding structured representations that neatly reflect sequence dynamics and stimulus identity after training. Finally, we found changes in the population-level temporal response patterns that reveal a reorganization of stimulus-driven activity that might reflect a coding strategy supporting temporal predictions.

When an expected transition was omitted by holding the preceding image on the screen, we found an elevated response after training that peaked when the omitted element would have been seen and was sustained throughout the omitted stimulus period. This contrasted with a decaying response observed at baseline. The temporal specificity of the response supports our previous findings that V1 responses are actively modulated by temporal expectations formed during exposure to spatiotemporal visual sequences (Price et al., 2023). Other work outside of our lab has provided evidence for omission-type prediction errors in L2/3 of V1, but the experimental paradigms vary in ways that make direct comparisons difficult. Fiser *et al*. 2016 found a subset of cells that responded anticipatorily to an omitted grating that was expected to be found at a particular location along a linear track. In this case, the task involved spatial location, locomotion, water reward, and perhaps most crucially, the complete omission of a grating at an expected location rather than a held-over grating and no predetermined timing of the grating presentations. In another study, Garrett *et al*. observed ramping activity in vasoactive intestinal peptide (VIP) expressing inhibitory cells but not excitatory cells in L2/3 in response to an omitted image (Garrett et al., 2020). The images were separated by 500 ms gray periods and omitted activity could have been predicated in part by visual transitions that were absent here and in Price *et al*. 2023.

When an expected grating was substituted with a different (but familiar) grating, we did not observe elevated activity. This finding contrasts with our earlier work in layer 4 and is inconsistent with predictive coding theory. We do not see any evidence of prediction errors in C-selective neurons. This negative result is mirrors recent work in monkey V1 and human EEG that found little evidence of substitution-type prediction errors in a conceptually similar experiment involving passive exposure to standard and deviant visual sequences (Solomon et al., 2021). This work recorded from electrodes positioned at unknown cortical depths. Our finding that substitution errors are found in deeper layers but absent superficially raises the possibility that the negative result in primates could reflect a mismatch between recorded lamina and the location of prediction error cells.

It is also possible that substitutions do generate error signals in mouse layer 2/3 but that the dynamics of calcium imaging (wither the dynamics of calcium signals or acquisition frame rate) are too slow to accurately capture rapid prediction error responses. This has been suggested as an explanation for why Stimulus-selective Response Potentiation (SRP), a similar but mechanistically distinct form of visual plasticity, is clear in electrophysiology but not in calcium signals (Montgomery et al., 2022). Recent work in the auditory cortex found that prediction error cells respond with rapid, transient responses lasting only about 40 ms (Audette and Schneider, 2023). Finally, we cannot rule out that exposure to the ACBD sequence during baseline prevented an error by durably encoding this sequence as “familiar”, though this seems unlikely given the structure of our training paradigm. Regardless, this finding is in relatively stark contrast to the large effect ordinal expectation exerts on in LFP recordings or spiking cells in layer 4 (Gavornik & Bear, 2014; Sidorov et al., 2020; Price et al., 2023). This discrepancy could reflect the fact that the LFP represents dendritic currents (e.g. inputs, see Buzsáki et al. 2012) rather than somatic spiking (e.g. outputs) and includes inhibitory activity that we are functionally blind to in these experiments. Additional work is required to resolve this issue.

Consistent with principles of efficient coding (Price & Gavornik 2022), we found that training had the effect of creating relatively sparse response patterns. Using two different measures, we estimate that the training reduced the number of visually modulated cells by 20- 30%. The sparsification may be related to the qualitative changes in principal component space, where training creates principal components with dynamics that neatly correspond to individual stimuli. In addition to being more efficient, a sparser orthogonalized code could also be easier for other brain regions to interpret. Admittedly, this argument would be stronger if element or temporal decoding accuracy had increased with training. While it is certainly true that brain processes operate very differently than a linear decoder and may take greater advantage of modified dynamics, it is also worth noting that the decoders on day 5 were as accurate as those trained against baseline data despite the smaller proportion of visually responsive neurons on day 5.

Neural sequences have been proposed as one of several mechanisms for representing durations, and there is growing support for this idea in different parts of the brain (Tsao et al., 2022). In this view, different cells fire preferentially at different delays relative to the onset of an external event, thereby forming a stable neural trajectory from which different points in time can be read out. Our data suggests that cells learn specific “time fields” (i.e., locations of peak activity) that, at the population level, uniquely represent timepoints spanning the 250 ms sequence element durations. By focusing our analysis on cells with consistently timed activity, we found significant differences in sequential activation following training. Before training, relatively few cells peaked at or near stimulus onset, and time field distributions became increasingly dense throughout the ∼200 ms post-onset window (figure 5B). After training, more cells peaked at stimulus onsets, and intermediate durations were less densely represented. Our analysis of time fields during the interstimulus gray period showed a surprising degree of temporal structure, though no evidence of experience-dependent plasticity. A cluster of peaks in the 300 and 750 ms ranges are tantalizingly close to the 250 ms element time and 800 ms gray period intervals, but the fact that they are in early traces from baseline recordings makes it unlikely to reflect a learned response. These peaks might be related to visually evoked theta-range oscillations reported previously (Gao et al., 2021; Kissinger et al., 2020; Levy et al., 2017; Zold & Hussain Shuler, 2015) though these are reported to develop with familiarity (Kissinger et al., 2018).

Our findings build on previous work to show that sequence plasticity modifies evoked responses in superficial layers of V1 as they do in thalamocortical input layers, albeit with notable differences. The development of sustained responses during element omissions appears to be a common feature of both L2/3 and L4, whereas we did not observe the emergence of substitution-type prediction errors in L2/3 as was previously observed in deeper layers. Overall, the work continues the recent tradition of providing ambiguous support for the idea that cortical dynamics are best described by predictive coding models while simultaneously demonstrating that evoked dynamics in V1 are far more complex than canonical visual processing models suggest.

## Methods

### Animal subjects

A total of eight mice (3 male, 5 female) aged 2-5 months were used for this study [CaMKII-tTA:tetO-GCaMP6s (Jackson Laboratories stock numbers 007004 and 024742)]. Mice were housed in a climate-controlled environment on a standard 12-hour light-dark cycle and were provided with food and water *ad libitum*. Cranial windows were implanted in mice at P40-P70. Experiments were performed during the mouse’s light cycle. All procedures were approved by the Institutional Animal Care and Use Committee (IACUC) of Boston University.

### Cranial Windows

Mice were briefly anesthetized with isoflurane (∼2% by volume in O2) and placed on a stereotactic surgical stage with warming pads to maintain body temperature. Anesthetic gas (1- 2%) was passively applied through a nose mask and adjusted as necessary to maintain respiratory rate and suppressed hind-paw and tail-pinch reflexes. Eyes were coated with a thin layer of Soothe eye lubricant (Bausch and Lomb, Canada). The scalp was shaved and opened with a rostrocaudal incision at the midline using scissors, and the periosteum was removed. A 5 mm circular craniotomy was made over the left visual cortex (2.9 mm lateral and 0.5 mm anterior to lambda). A 5 mm cranial window (Deckglaser) was placed over the craniotomy and secured with Metabond. A steel headplate was then affixed to the skull. Mice recovered for 7-10 days prior to imaging.

### Retinotopic Mapping

1-3 days prior to beginning the sequence learning experiment, a retinotopic map of the visual cortex was generated using widefield one-photon microscopy. The brain was illuminated with a 470 nm light source (X-Cite 200DC), and images were acquired through a 10x objective, a sCMOS camera (Thorlabs Quantalux, CS2100M-SB), and ThorImageLS 3.0 (Thorlabs Inc.). A 22-inch LED monitor (1920 x 1080 pixels, refresh rate 60 Hz) was positioned 15 cm from the mouse’s right eye, and a narrow sweeping stimulus moved across the screen in four directions (left-to-right, right-to-left, top-to-bottom, and bottom-to-top). Trial-averaged responses were used to determine the location of binocular V1.

### Two-photon Imaging

Two-photon Ca2+ imaging was performed on days 1 and 5 using a Bergamo microscope (Thorlabs Inc., Newton, NJ, USA) controlled by ThorImage OCT software (ThorImageLS, v3). The visual cortex was illuminated with a Ti:Sapphire fs-pulsed laser (Mai Tai Deep-See, Spectraphysics) tuned to 920 nm. The laser was focused onto L2/3 of binocular V1 through a 16x water-immersion objective lens (0.8NA, Nikon). Ca2+ transients were obtained from neuronal populations at a resolution of 512 x 512 pixels (sampling rate ∼30 Hz). The obtained images were motion-corrected using CaImAn (Giovannucci et al., 2019). Segmentation, neuropil subtractions, and deconvolution were performed using Suite2p (Pachitariu et al., 2016).

### Visual Stimulus

Visual stimuli were generated and displayed using MATLAB with the PsychToolbox extension (Brainard, 1997), with custom software (https://github.com/jeffgavornik/VEPStimulusSuite) used to control timing and hardware signals. Stimuli were displayed on a 22-inch LED monitor (1920 x 1080 pixels, refresh rate 60 Hz) positioned 25 cm directly in front of the mouse to stimulate binocular V1. All stimuli were matched for luminance.

Four images (referred to as A, B, C, and D), each an oriented sinusoidal grating, had angles 15, 75, 165, and 120 degrees, respectively) with a spatial frequency of 0.05 cycles/degree. These images were displayed for 250 ms each. Images were combined into three sequences (ABCD, ABBD, and ACBD). Every sequence presentation was followed by an 800 ms period of gray screen. Therefore, each sequence presentation, including its following gray period, lasted 1800 ms. Note: since there were no gray periods between images in a sequence, BB in ABBD was essentially a single 500 ms presentation of B.

### Experimental Design

On day 0, all sequences (ABCD, ABBD, and ACBD) were shown in blocks of 100 sequences (e.g. ABCD x 100, ABBD x 100, ACBD x 100), and each block was separated by a 10-second rest period (see Figure 1). This structure was repeated 5 times, yielding 500 presentations of each sequence on any given day. Day 0 and day 1 were separated by a two-day buffer period. During the training period (days 1-4), only ABCD was shown and in 5 blocks of 100 presentations. Day 5 (testing) was identical to day 0. Additionally, there was a 1-minute gray period preceding and following the experiment.

### Stimulus and context selectivity

For each cell, we computed trial- and time-averaged responses to all stimuli and gray periods after applying a 67 ms offset from stimulus onset to account for the information delay from retina to L2/3 (time windows were still 250 ms for images and 800 ms for gray periods after applying the offset). The duration of this offset/delay was verified by looking at individual cell responses after stimulus onset. We then compared the mean activity for a given stimulus with the mean activity over all other stimuli. If the mean activity for a given stimulus was over two standard deviations of the other stimuli, it was assigned selectivity for that stimulus. This was first done for stimuli regardless of sequence context. The procedure was performed again with sequence context taken into account to look for cells that were primarily active within a particular sequence (e.g., cells that fired to image C in ACBD but not ABCD). Manual inspection of all cells (n=2868) validated this approach for >80% of cells. Manual curation was used to correct algorithmic classifications in a small number of cases (22 cells) showing mixed selectivity.

### Prediction error (PE) Ratios

PE ratios were computed by extracting the trial- and time-averaged (deconvolved) activity for each cell given an unexpected/deviant element and an expected/standard element. Note that a two-acquisition frame offset (66 ms) was applied to account for the information delay from retina to L2/3. The ratio is computed by dividing the averaged activity during the deviant element by averaged activity during the standard element. For omission-type PE ratios, only B-responsive cells were considered, and A**B**BD and AB**B**D were used as the deviant and standard elements, respectively, in the main text. An equivalent analysis was performed using A**B**CD as the standard element in supplementary figure 2. For substitution-type PE ratios, only C-responsive cells were considered, and ACBD and ABCD were used as the deviant and standard elements, respectively.

### Principal Component Analysis

Trial-averaged responses for ABCD, ABBD, and ACBD were computed and pooled across mice. These responses were then concatenated across time yielding a time-by-cell matrix. After performing PCA, the resulting principal components were then split in time to show how the shared set of principal components behaved during each of the three sequences.

### Sparseness estimation

We estimated sparseness of single-cell responses in two ways.

1. Stimulus selectivity: We counted the number of neurons that are driven by a particular stimulus more than two standard deviations above their mean rates. For each neuron, we computed the trial- and time-averaged responses for each image. The mean firing rate and standard deviation were computed across time regardless of stimulus. A neuron was considered as responsive to a given stimulus if its response was more than two times this standard deviation plus its mean firing rate.
2. Visual modulation: We counted the number of cells that had significantly different activity during gratings vs gray periods. For each neuron, we collected and time-averaged 266 ms chunks for gratings and gray periods separately. We then performed a KS-test on these two groups, and if the two distributions were significantly different (p < 0.05), then the cell was classified as visually modulated. Note 266 ms chunks were separated in time by a 133 ms gap in order to mitigate correlations between temporally neighboring samples.

### Correlation analysis

For each sequence presentation, we extracted time-averaged responses for each stimulus (excluding the first 66 ms after image onset). We then computed Pearson-correlation coefficients for all 6 pairs in the presentation. For example, for the first presentation of ABCD, we would first extract and time-average population vectors for each image A, B, C, and D. We would then compute Pearson correlation coefficients between all 6 pairs: A to B, A to C, A to D, etc. In this way, we had 500 presentations x 6 pairs = 3000 coefficients for each sequence for each day. We compared coefficients between days with KS-tests.

### Stimulus decoding

Time-averaged responses to all stimuli were used to train a linear decoder. We excluded the first two time bins (67 ms) to account for the information delay to L2/3 and avoid contamination from the previous stimulus. Data was randomly split into two equally sized groups (250 trials each) for training and testing. This was repeated 100 times. Accuracy by random chance was 1/15 = 6.7%.

### Block decoding

For each stimulus, time-averaged responses were computed for all cells and all trials (excluding the first two time bins as above). A support vector classifier was then trained on 50 randomly chosen trials from each block and tested on the remaining trials (n=250 trials for each group).

This was repeated for all 12 sequence elements separately (**A**BCD, **A**BBD, **A**CBD, …), and the results were pooled and averaged to produce the confusion matrix shown in figure 4B and supplementary figure 5.

### Temporal decoding

Data from all intersequence gray periods (n=1500) was collected and randomly split into two equally sized groups (750 trials each). The decoder was trained on one set and tested on the other. This was repeated 100 times. Accuracy by random chance was 1/24 = 4.2%.

### Time field estimation

Trials were split into even/odd groups, and responses were averaged over trials for both groups. For each cell, we found where the cell fired maximally in each group. If the location of the even- group maximum was within 133 ms (4 time bins) of the odd-group maximum, the cell was considered to have a temporally consistent firing pattern (∼60% of cells on both days), and the remaining cells were discarded. Consistently firing cells were assigned a time bin based on their points of maximum activity using data averaged over all trials.

### Code/Software

All data analysis was performed in Python using the Python scientific stack (Numpy, Scipy, Sci- Kit Learn). For decoder analysis, we used sci-kit learn’s support vector classifier with a linear kernel. Analysis code is published in a Jupyter notebook available at https://gavorniklab.bu.edu/supplemental-materials.html.

### Statistics

We used the nonparametric KS test to compare distributions throughout the paper. To confirm that the hierarchical structure of our experiment was not impacting our conclusions, we also applied a nonparametric bootstrap resampling method for multilevel data (Saravanan et al., 2020) to estimate mean traces over stimulus-selective cell groups and their respective mean PE ratios.

Each iteration of the bootstrap (1000 in all) follows the same basic procedure: randomly resample mice (with replacement), combine data from this subset into a single pool, resample from this pool (with replacement), and then take and store the mean of the samples. We used this resampled distribution to calculate the median population response with 95% confidence intervals and PE ratio statistic as shown in supplemental figure 8 (additional methodological details included in supplemental materials).

## Author Contributions

Experimental conception and design: SGK and JPG. Experiments and surgeries: SGK, CM. Data analysis and figures: GSK and CM. Writing: SGK and JPG.

## Data Availability

All data and analysis code is available at https://gavorniklab.bu.edu/supplemental-materials.html.

## Acknowledgements

Special thanks to Marc Howard for his insight and Byron Price for his feedback and discussions throughout the design and analysis of these experiments. This work was supported by NEI R001EY030200.

