## supplemental materials for "Learned response dynamics reflect stimulus timing and encode temporal expectation violations in superficial layers of mouse V1"

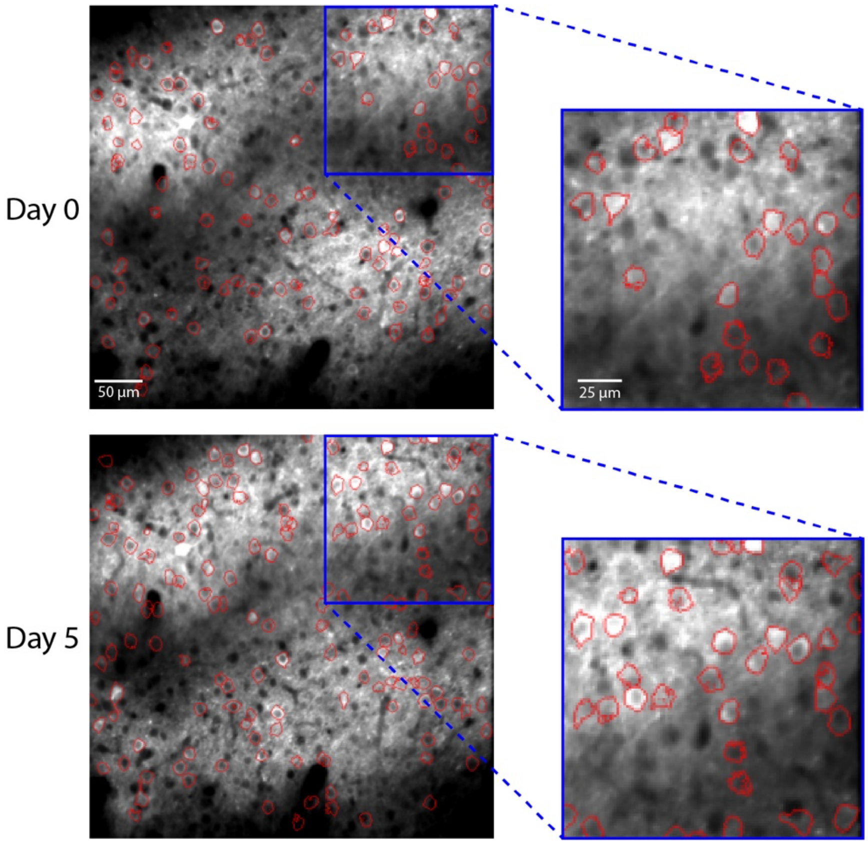


**Supplementary Figure 1: Fields-of-view across days.** Representative field-of-view (FOV) for one mouse on day 0 (upper panels) and day 5 (lower panels). ROI boundaries are drawn in red and laid over mean intensity images. Right panels display magnified subregions. Cells were identified based on activity, and all identified cells were analyzed on both days.

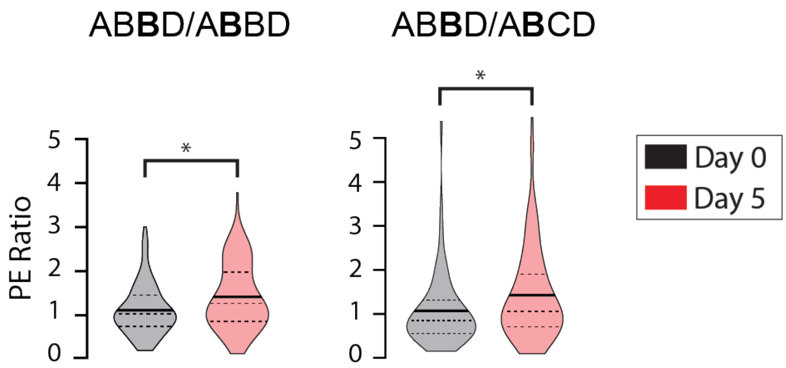


**Supplementary Figure 2: Omission-type prediction-errors.** Omission-type PE ratios can be computed using activity during AB**B**D against either A**B**BD (left panel) or A**B**CD (right panel). The left panel (also shown in the main text), which uses A**B**BD as the standard element, shows the distributions of PE ratios to be significantly different between days 0 and 5 ($p << 0.05$; n=244; KS-test). On day 0, mean PE=1.10 (n=137). On day 5, mean PE=1.44 (n=107). Conclusions from this analysis do not change if it is repeated using A**B**CD as the standard element for the PE calculation. As shown in the right panel, there is still a statistically significant differences between days 0 and 5 ($p << 0.05$; n=244; KS-test). On day 0, mean PE=1.04 (n=137). On day 5, mean PE=1.43 (n=107). Dashed horizontal lines indicate the 25$^{th}$, 50$^{th}$, and 75$^{th}$ percentile. The solid horizontal lines indicate population means.

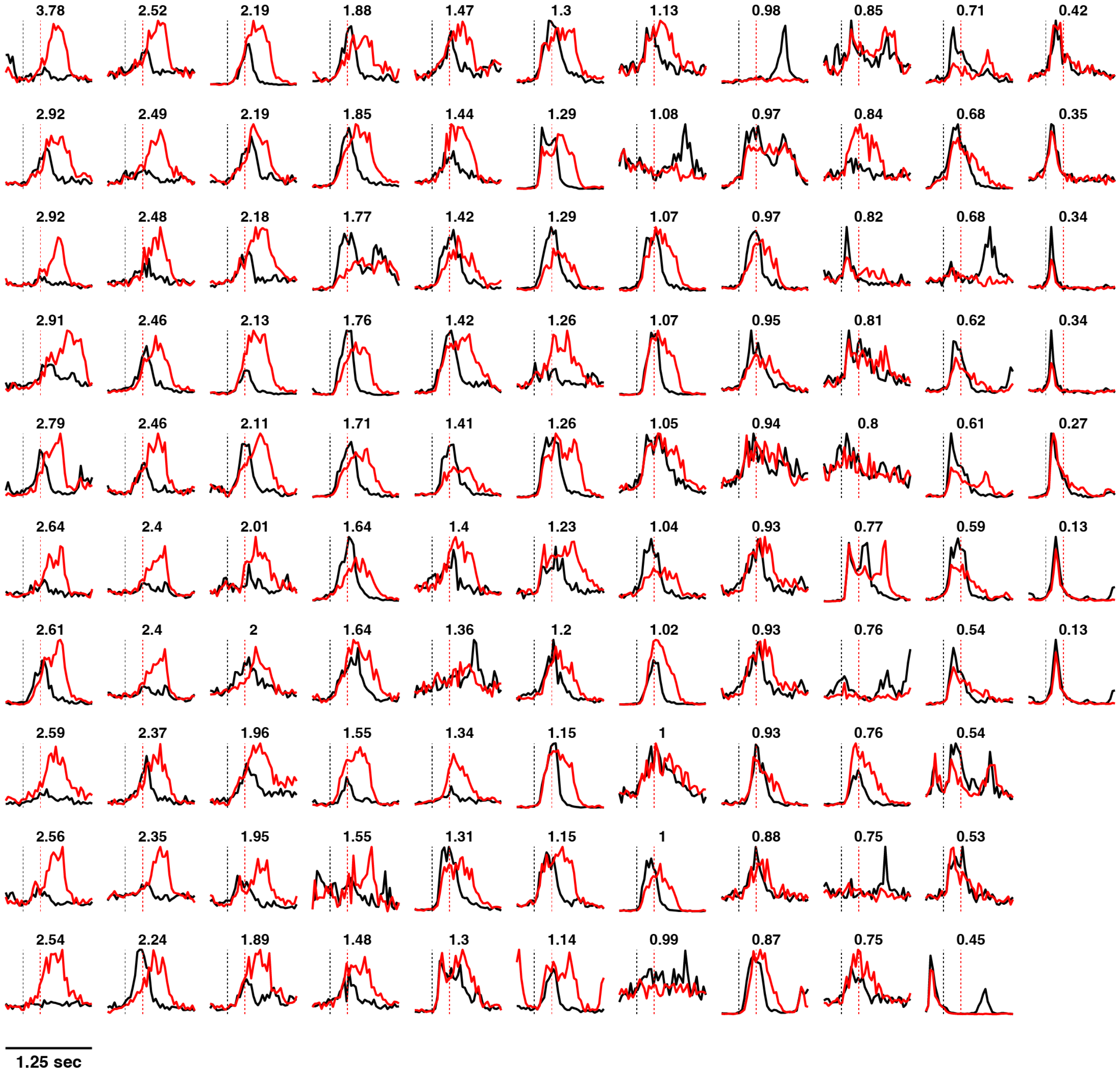


**Supplementary Figure 3: B-responsive cell traces.** Trial-averaged responses for all 107 B-responsive cells during ABCD (black) and ABBD (red) on day 5. The black dashed line indicates onset of the first B (A**B**CD or A**B**BD), and the red dashed line indicates the onset of the following element (AB**C**D or AB**B**D). Plots are ordered from top-to-bottom, left-to-right and in descending order of PE ratio (PE ratios are overlayed on plots). PE ratios were computed using A**B**BD and AB**B**D as the standard and deviant elements, respectively. Cells with high PE ratios typically respond to the first B in the sequence (A**B**BD), and they continue firing until peaking within AB**B**D. Only 1 cell (top-left) fires uniquely to the omitted element.

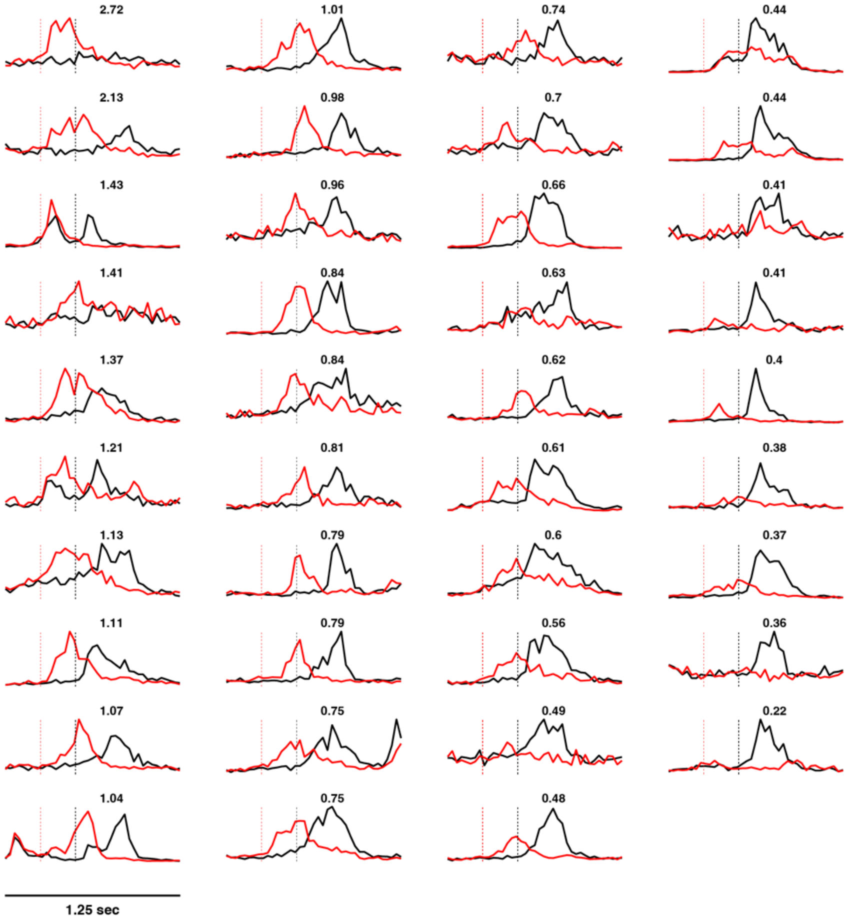


**Supplementary Figure 4: C-responsive cell traces.** Trial-averaged responses for all 39 C-responsive cells during ABCD (black) and ACBD (red) on day 5. Dashed red lines indicate the onset time of the second element in the sequence (A**B**CD or A**C**BD), and the black dashed line indicates the onset of the third element (AB**C**D or AC**B**D). Plots are ordered from top-to-bottom, left-to-right and in descending order of PE ratio (PE ratios are overlayed on plots). PE ratios were computed using AB**C**D and A**C**BD as the standard and deviant elements, respectively. All cells with PE ratios > 1 responded strongly to A**C**BD relative to AB**C**D, though only one (top-left) responded exclusively to A**C**BD.

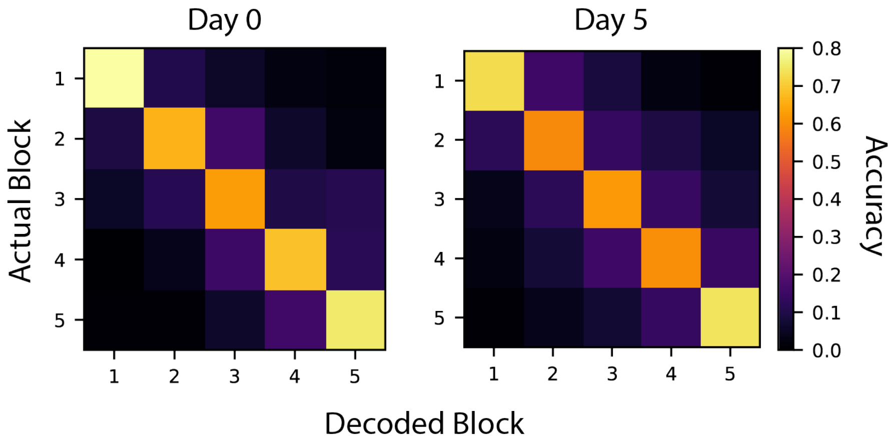


**Supplementary Figure 5: Block decoding with normalized population vectors.** In principle, slow changes in overall firing rates due to photobleaching, arousal, etc. could account for a decoder's ability to accurately predict which block neural responses belong to. To test whether a simple rescaling process can account the block decoding results shown in figure 4B, we repeated our analysis using normalized population vectors. Similar to the findings reported in the main text, decoders accurately predict which block responses came from. Responses from nearby blocks were more likely to be misclassified compared with responses from distant blocks. Decoder accuracy was 70% on day 0 and 65% on day 5. Apart from normalization, the same training procedure was used here as in the main text.

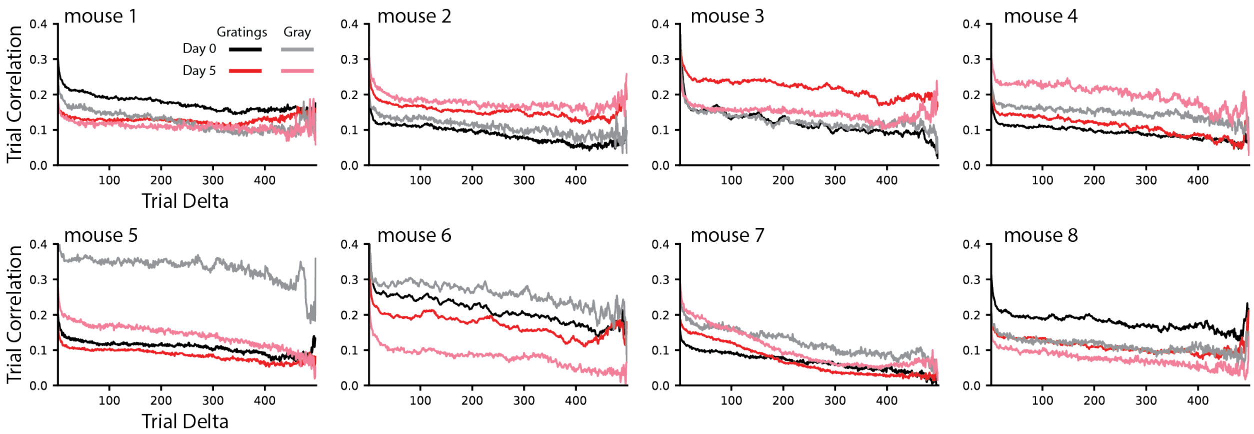


**Supplementary Figure 6: Representational drift for individual mice.** Pearson-correlation coefficients were computed between all pairs of same-stimulus response vectors and grouped by distance (in time/trial) between pairs for mice individually. The number of pairs per grouping ranges from 499 (when trial delta = 1) and 2 (when trial delta = 499), and the group averages form each line. Responses to nearby trials were more correlated/similar than distant trials in all conditions and mice.

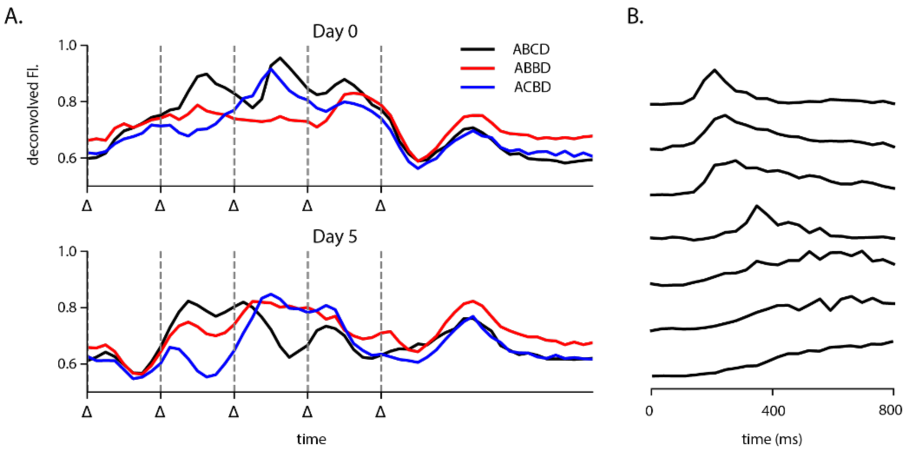


**Supplementary Figure 7: Mean traces and selected single-cell responses during gray periods.** (A) Trial- and population-averaged activity for all cells and sequences on day 0 (top) and day 5 (bottom). Triangles on the x-axis mark stimulus changes which occur every 250 ms except for the trailing gray period which lasts 800 ms. The population-averaged activity increases ~300 ms after gray onset described in figure 6 of the main text. (B) Trial-averaged activity for a selected subset of cells during the gray period only on day 0. Select cells exemplify the ~300 ms activity increase and the late ramping activity (see figure 6A heatmaps showing activity of all cells).

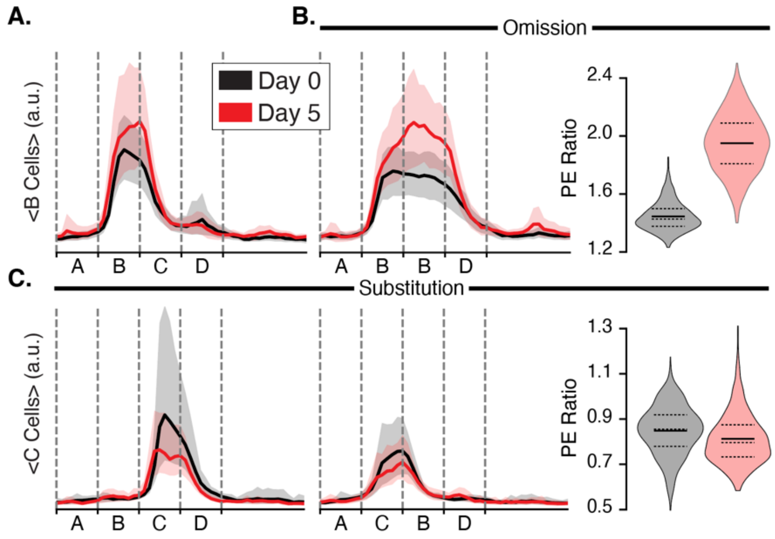


**Supplementary Figure 8: Hierarchical bootstrapped statistics.** Mean responses estimated from bootstrapped distributions for B and C responsive cells match responses in Figure 2. **A**. Bootstrapped mean responses of B-responsive cells to ABCD on days 0 (black) and 5 (red). Solid lines show the mean response from the resampled distributions, shaded regions show 95% confidence intervals. **B.** Bootstrapped responses of B-responsive cells to ABBD (left) and distribution of mean omission-type PE ratios (right). Mean PE ratios are significantly larger following an omission on day 5 (mean PE=1.95) relative to day 0 (mean PE=1.45)(p=<<0.05; n=1000; KS-test) **C.** The response of C-responsive neurons to ABCD and ACBD show a slightly smaller PE ration following substitution on day 5 (mean PE=0.81) compared with day 0 (mean PE=0.85)(p<<0.05; n=1000; KS-test). These results, which account for the hierarchical nature of our experimental design, confirm the conclusions in the main body.

To illustrate how the bootstrap was used to estimate mean traces and PE ratios, we will use day 0 omission-type responses as an example. In each iteration, 8 mice were randomly selected (with replacement) from the original group of 8. Trial-averaged day 0 ABBD response traces for B-responsive cells in the resampled subset were then pooled into a single matrix, where each row represented the trial-averaged activity for a cell. We selected rows from this matrix (with replacement) equal to the total number of rows in the pooled matrix and calculated and stored the mean trace and PE over these samples. After 1000 iterations, 95% confidence intervals were generated by finding the 2.5th and 97.5th percentile of the bootstrapped mean traces and PE ratios. Violin plots were generated to show the distribution of PE ratios on days 0 and 5. Unlike in figure 2D, where each sample is a PE ratio for a single cell, samples in these distributions represent the mean PE ratios computed over the entire resampled population of cells from a single bootstrap iteration.
